# COMPLETE GENOME SEQUENCE OF *VIBRIO SYNGNATHI SP. NOV*., A FISH PATHOGEN, ISOLATED FROM THE KIEL FJORD

**DOI:** 10.1101/2023.04.21.537764

**Authors:** Cynthia Maria Chibani, Robert Hertel, Meina Neumann-Schaal, Henry Goehlich, Kim Wagner, Boyke Bunk, Cathrin Spröer, Jörg Overmann, Michael Hoppert, Mareike Marten, Olivia Roth, Heiko Liesegang, Carolin C. Wendling

**Affiliations:** Georg-August University Goettingen, Institute of Microbiology and Genetics, 37077 Goettingen, Germany; Institute for General Microbiology, Kiel University, 24118 Kiel, Germany; Leibniz Institute DSMZ-German Collection of Microorganisms and Cell Cultures, Inhoffenstr. 7B, 38124 Braunschweig, Germany; GEOMAR, Helmholtz Centre for Ocean Research, Marine Evolutionary Ecology, Düsternbrooker Weg 20, 24105 Kiel, Germany; Marine Evolutionary Biology, Zoological Institute, Kiel University, 24118 Kiel, Germany; ETH Zürich, Institute of Integrative Biology, Universitätsstraße 16, 8092 Zürich, Switzerland

**Keywords:** *Vibrio*, Genomic comparison, Fish pathogen, pan-genome, virulence factors, *Vibrio Splendidus* clade, *Vibrio*, complete genome sequence

## Abstract

A new *Vibrio* strain K08M4^T^ was isolated from the broad-nosed pipefish *Syngnathus typhle* in the Kiel Fjord. Infection experiments revealed that K08M4^T^ is highly virulent for juvenile pipefish. Cells of strain K08M4^T^ are Gram-stain-negative, curved rod-shaped and motile by means of a single polar flagellum. The strain can grow aerobically at 9 to 40°C, at pH 4 to 10.5 and tolerates up to 12% (w/v) NaCl. The most prevalent (> 10%) cellular fatty acids of K08M4^T^ were C_16:1_ *ω*7*c* and C_16:0._ Whole-genome comparisons revealed that K08M4^T^ represents a separate evolutionary lineage which is distinct from other *Vibrio* species and falls within the *Vibrio Splendidus* clade. The genome is 4,886,292 bp in size, consists of two circular chromosomes (3,298,328 bp, 1,587,964 bp), and comprises 4,178 protein-coding genes and 175 RNA genes. In this study, we describe the phenotypic features of the new isolate and present the annotation and analysis of its complete genome sequence. Based on these data, the new isolate represents a new species for which we propose the name *Vibrio syngnathi*. The type strain is K08M4^T^ (=DSM 109818^T^).

**Supplementary material:** One supplementary figure and six supplementary table are available with the online version of the Manuscript.

## Full Text

The genus *Vibrio* comprises 137 validly published species (https://lpsn.dsmz.de/genus/vibrio) which have been grouped into 16 monophyletic clades [1, 2]. The *Vibrio* S*plendidus* clade, which contains 17 closely related *Vibrio* species constitutes the largest clade within the genus *Vibrio [3]*. The phenotypic variability of additional species, which did not fit exactly the description of *V. splendidus* led to the proposal of the term *Vibrio splendidus*-like, which is now widely used for isolates that cannot unequivocally be assigned to the species *V. splendidus*.

Bacteria of the *V. Splendidus* clade are dominant members of the costal bacterioplankton, in sediments, and various marine organisms, for a review see [4]. Various *V. splendidus*-like strains have been found to cause vibriosis. Vibriosis is one of the most prevalent diseases affecting marine life accounting for major economic losses in aquaculture [4, 5]. Vibriosis causes high rates of mortality in aquaculture animals such as turbot, scallops, clams, and oysters [6-9] and can even spread to humans through the consumption of infected seafood [10, 11]. Virulence of these strains is multifactorial and regulated by genetic determinants, e.g. prophages [12] or pathogenicity islands [13] as well as extrinsic factors, such as temperature [14] or salinity [15]. Furthermore, a recent epidemiological survey of the *V*. S*plendidus* clade suggests an unexplored diversity of virulence factors and a massive lateral transfer of virulence genes within this clade [16].

Despite a significant body of empirical research, our understanding of the factors that contribute to the virulence of *V. splendidus-*like strains is far from being complete. Whole-genome sequencing can provide further insights into potential virulence mechanisms of this pathogen. At the time of writing, 819 complete *Vibrio* genome sequences and 13 *V. splendidus*-like strains were publicly available (NCBI, Date of access 22.05.2022).

In this study, we sequenced and annotated the complete genome of strain K08M4^T^ which was found to represent a new *Vibrio* species, *Vibrio syngnathi*. We follow the proposed minimal standards for the use of genome sequence data for taxonomic purposes [17] and provide first insight into the genomic basis of its pathogenicity.

### Origin and isolation

Strain K08M4^T^ was isolated during a previous study from the intestine of a healthy adult pipefish collected in the Kiel Fjord in 2010 [for a detailed description see [18]] and was shown to be pathogenic to juvenile pipefish in laboratory experiments. Based on a multi-locus sequencing approach, K08M4^T^ was initially affiliated to the *V. Splendidus* clade [19]. The whole-genome comparison in the present study supports the consideration that K08M4^T^ represents a distinct species within the *V. Splendidus* clade.

### Phenotypic characterization

K08M4^T^ was routinely cultivated on *Vibrio*-selective thiosulfate-citrate-bile-sucrose (TCBS) agar plates or in liquid culture using Medium101 (Medium101: 0.5% (w/v) peptone, 0.3% (w/v) meat extract, 1.5% (w/v) NaCl in MilliQ water) at 25°C.

Cell morphology of strain K08M4^T^ was determined by electron microscopy of negatively stained cells using a Jeol 1011 transmission electron microscope (Eching, Munich, Germany) as described in [20]. Briefly, cells were grown in liquid Medium101 for 12 h overnight at 25° C with constant shaking at 230 rpm as pre-culture. The main culture was inoculated with 1.5% v/v grown for 3 hours and adsorbed onto Formvar-carbon grids (Plano, Wetzlar, Germany) prior to staining. Phosphotungstic acid, dissolved in pure water (1% resp 0.5% *w*/*v* final concentration) and adjusted to pH 7 with sodium hydroxide served as a staining solution. Images were captured using a Gatan Orius SC 1000A camera and processed with the Gatan Digital Micrograph software package (vs. 1.84.1177 Pleasanton, CA, USA). Cells of strain K08M4^T^ are slightly curved (1.5-2.5 μm long and 0.5-1 μm wide) with a monotrichous flagellum (Figure 1).

**Figure 1:**
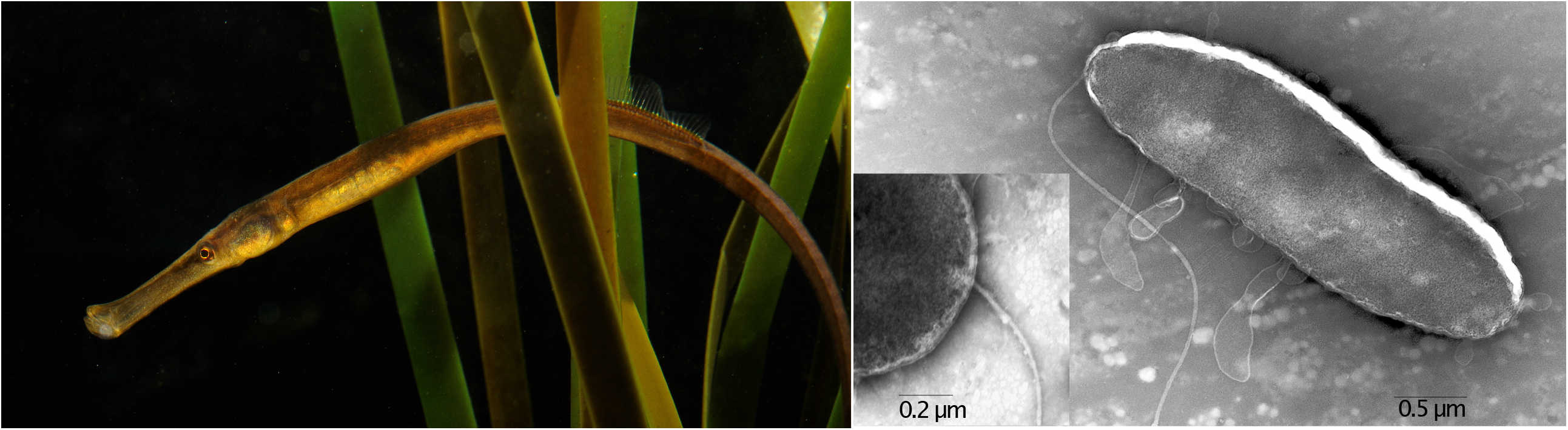
Left: Broad-nosed pipefish, *Syngnathus typhli* (by Uli Kunz), from which K08M4^T^ was isolated from; right: Transmission electron microscope image of K08M4^T^ after 3 hours of growth in medium 101.

Salinity and temperature ranges were determined by generating growth curves over 24 h-time intervals of 1:100 dilutions of overnight cultures using a microplate reader (TECAN infinite M200) at different NaCl concentrations ranging from 0-15 % (w/v) at 25° C with 0.5 % intervals, as well as across a temperature gradient of 4 to 45° C with 5° C intervals at 1.5% (w/v) NaCl and different pH levels ranging from pH 3.5 to 11 at intervals of 1.0 pH unit. Strain K08M4^T^ produces yellow-colored, round colonies on TCBS agar plates and cream-colored, round colonies on Medium101 plates after incubation for 24h at 25° C. The strain did not grow below 9° C and showed optimum growth between 25 and 35° C at 0.4 to 12 % NaCl, with optimum growth occurring at 1.5 % NaCl and between pH 4 and 10.5.

We performed a comparative phenotypic analysis between strain K08M4^T^ and three closely related species obtained from the German Collection of Microorganisms and Cell Cultures DSMZ: *V. tasmaniensis* DSM 17182^T^, *V. splendidus* DSM 19640^T^, and *V. splendidus* DSM 26178^T^. The Gram reaction was determined by KOH and aminopeptidase using the Bactident Aminopeptidase (Merck, 113301) tests [21]. Oxidase activity was tested by the method of Kovacs et al. (1956) [22]. Catalase tests were performed by mixing freshly grown bacterial cells with 10% H_2_O_2_, followed by examining gas bubble formation. Chitinase activity was confirmed using the Chitinase Assay Kit (Sigma Aldrich, CS0980). Activities of constitutive enzymes and other physiological properties were determined using API 20E, API ZYM and API 50CH (bioMérieux) kits according to the instructions of the manufacturer. Susceptibility to the vibriostatic agent 2,4-diamino-6,7-diisopropyl pteridine phosphate salt (O/129, Sigma) was determined by means of discs containing 150μg O/129. Fatty acid analysis was performed using the standard protocol of the Sherlock Microbial Identification (System package version 6.4 with the TSBA6.0 database), according to the technical instructions provided by the manufacturer [23]. Combined analysis by gas chromatography coupled to a mass spectrometer was used to confirm the identity of the fatty acids based on retention time and mass spectral data [24]. Major characteristics and unique phenotypic characteristics differentiating K08M4^T^ from its close relatives *V. tasmaniensis* DSM 17182^T^, *V. splendidus* DSM 19640^T^, and *V. splendidus* DSM 26178^T^ are summarized in Table 1, full results are given in Table S1-S3. The cellular fatty acid profile of K08M4^T^ in comparison to *V. tasmaniensis* DSM 17182^T^ and *V. splendidus* DSM 19640^T^ is listed in Table 2, full results for strain K08M4^T^ only, are given in Table S4.

**Table 1:** Differential characteristics between strains K08M4^T^ and the reference strains of phylogenetically related species Strains: 1. DSM 109818^T^ (this study), 2. *V. tasmaniensis* DSM 17182^T^ (this study), 3. *V. splendidus* DSM 19640^T^ (this study), 4 *V. splendidus* DSM 26178^T^ (this study), 5. *V. pomeroyi* LMG 20537^T^ [43], 6 *V. gallaecicus* LMG 24045^T^ [44, 45], 7 *V. cholerae* ATCC 14035^T^ [46]; +, positive reaction; -, negative reaction; ND, no data available

|  | 1 | 2 | 3 | 4 | 5 | 6 | 7 |
| --- | --- | --- | --- | --- | --- | --- | --- |
| ADH | - | + | + | + | + | - | - |
| VP | - | - | - | - | - | - | ± |
| ONPG | - | - | + | + | + | - | ND |
| O/129 susceptibility | + | - | + | - | + | + | - |
| Gelatin hydrolysis | - | - | + | + | + | + |  |
| Acid from |  |  |  |  |  |  |  |
| Galactose | - | - | + | + | ND | ND | + |
| Mannitol | - | + | + | + | + | + | + |
| Amygdalin | + | + | + | + | + | - | ND |
| Melibiose | - | - | + | + | - | - | - |
| Sucrose | + | - | - | - | + | - | + |
| 2-Ketogluconate | - | - | + | - | ND | ND | - |
| Mannose | - | + | + | + | ND | ND | + |

A full Table containing all results from API ZYM, API 20E, and API 50CHE can be found in the supplementary material Table S1-S3.
ADH Arginin dihydrolase, VP Voges Proskauer, ONPG β-Galactosidase

**Table 2:** Fatty acid profile of strain K08M4^T^ DSM 109818^T^ in comparison to *V. tasmaniensis* DSM 17182^T^, 3. *V. splendidus* DSM 19640^T^. All data in the table are from the present study. Fatty acids present in amounts lower than 1% are not shown. Quantification by GC-FID, identity was confirmed by mass spectrometry.

| Fatty acid | K08M4<sup>T</sup> | *V. tasmaniensis*<br>DSM 17182<sup>T</sup> | *V. splendidus*<br>DSM 19640<sup>T</sup> |
| --- | --- | --- | --- |
| C<sub>12:0</sub> | 3.9% | 4.0% | 5.0% |
| C<sub>12:0</sub> 3OH | 2.7% | 2.4% | 2.5% |
| C<sub>14:0</sub> | 3.8% | 4.1% | 5.7% |
| C<sub>14:0</sub> 3OH | 1.8% | 1.7% | 1.9% |
| C<sub>16:0</sub> iso | 7.6% | 3.4% | 3.2% |
| C<sub>16:1</sub> ω7c | 39.9% | 43.8% | 40.7% |
| C<sub>16:1</sub> ω7t | 3.3% | 4.3% | 5.3% |
| C<sub>16:0</sub> | 19.2% | 21.6% | 19.4% |
| C<sub>18:1</sub> ω7c | 8.7% | 7.0% | 11.1% |
| C<sub>18:0</sub> | 1.0% | 1.1% | 0.5% |

A full table can be found in the supplementary material, Table S4.

### Genomic characterization

The Qiagen Genomic Tip/100 G (Qiagen, Hilden, Germany) was used to extract high molecular weight DNA from cells of strain K08M4^T^. The extracted DNA was then used to generate a SMRTbell™ template library for SMRTbell™, according to the instructions from Pacific Biosciences, Menlo Park, CA, USA, following the Procedure & Checklist -10 kb Template Preparation Using BluePippin™ Size-Selection System.

SMRT genome sequencing was carried out on the PacBio *RSII* (Pacific Biosciences, Menlo Park, CA, USA), taking one 240-min movie for a single SMRT cell using P6 chemistry at the Leibniz Institute DSMZ-German Collection of Microorganisms and Cell Cultures in Braunschweig. Genome finishing and annotation were performed at the Institute for Microbiology and Genetics at the Georg-August University of Göttingen.

We obtained a total of 64,464 post-filtered reads with a mean read length of 13,396 bp representing a genome coverage of 141.45-fold. Genome assembly was performed with the RS_HGAP_Assembly.3 protocol included in SMRT Portal version 2.3.0. The assembly resulted in two large contigs that could be associated with chromosomes 1 and 2. The genome has a size of 4,886,292 bp (chromosome 1: 3.2 Mb, chromosome 2: 1.5 Mb) and the G+C content of strain K08M4 is 44.06 mol%. The completeness and contamination levels of strain K08M4^T^ were computed using CheckM [25]. Based on the assembly of the genome without gaps or ambiguities, the completeness level (99.69%) and the contamination level (1.69%) of both chromosomes, a metagenome assembled genome would be labeled as “Finished” [26]. The genome project has been deposited at the European Nucleotide Archive under the ID number 345286 and accession number PRJNA345286. The accession number for chromosome 1 is CP017916 and for chromosome 2 is CP017917. The raw reads were deposited under the BioProject accession number PRJNA345286.

Automatic annotation, gene prediction, and rRNA and tRNA genes identification were carried out with Prokka (v1.11) [27]. Functional annotations were done by searching against InterPro [28], and COG databases [29]. A total of 4,353 genes were predicted, of which 4,178 encode proteins, 175 encode rRNAs, 43 encode tRNAs, and at least 46 encode ncRNAs. Twelve pseudogenes were found, with eight located on chromosome 1 and four located on chromosome 2. Of the predicted CDS, a functional prediction was made for 77.1 %. A COG prediction could be retrieved for 70.0%, with the remaining annotated as hypothetical proteins. We found a total of 148 insertion elements: 72 on chromosome 1 and 76 on chromosome 2. A putative 48.2-kb intact prophage was identified on chromosome 2 using PHASTER [30] and this region is predicted to encode 45 proteins, of which most are phage-related, whereas 23 are hypothetical or uncharacterized. In addition, one incomplete prophage was found on chromosome 2.

Whole genome based taxonomic analysis was done using the Type (Strain) Genome Server (TYGS) [31], a free bioinformatics platform provided by the DSMZ (https://tygs.dsmz.de) following the standard pipeline including recent updates [32]. The 11 closest type strain genomes were determined following the standard pipeline.

For phylogenomic inference, all pairwise comparisons among the set of genomes were conducted using GBDP and accurate intergenomic distances inferred under the algorithm ‘trimming’ and distance formula *d*_*5*_ [33]. 100 distance replicates were calculated each. Digital DDH values and confidence intervals were calculated using the recommended settings of the GGDC 3.0.

The resulting intergenomic distances were then used to infer a balanced minimum evolution tree with branch support via FASTME 2.1.6.1 including SPR postprocessing [34]. Branch support was inferred from 100 pseudo-bootstrap replicates each. The tree was rooted at the midpoint [35] and visualized using FigTree version 1.4.2 (http://tree.bio.ed.ac.uk/software/figtree/).

The type-based species clustering using a 70% dDDH radius around each of the 11 type strains was done as previously described [31]. Subspecies clustering was done using a 79% dDDH threshold as previously introduced [36]. The clustering yielded 12 species clusters and the provided query strains were assigned to 1 of these, K08M4^T^ was located in 1 of 12 subspecies clusters (Figure 2). The dDDH values between the complete genome sequence of strain K08M4^T^ and the genome sequences of the 11 most closely related species from the *V. Splendidus* clade are significantly below the species delination cut-off of 70% (Table 3).

**Table 3:** GGDC-Distance calculation for *V. syngnathi* K08M4T using the GGDC-server of DSMZ. Distances have been calculated using formulae 4, for details see [32, 33]– T for type strains.

| Species | DDH | Model C.I. | Distance | G+C difference | Accession Nr. Chr. 1 |
| --- | --- | --- | --- | --- | --- |
| *V. syngnathi* K08M4<sup>T</sup> | - | - | - | 44.06 mol% | CP017916.1 |
| *V. tasmaniensis* LMG 20012<sup>T</sup> | 58.9 | [56 - 61.6%] | 0.0536 | 0.03 | NZ_AP025510.1 |
| *V. atlanticus* LMG 24300 | 54 | [51.3 - 56.7%] | 0.0629 | 0.15 | NC_011753.2 |
| *V. atlanticus* CECT 7223<sup>T</sup> | 54.1 | [51.4 - 56.8%] | 0.0628 | 0.09 | NZ_AP025460.1 |
| *V. cyclitrophicus* ED008 | 39.2 | [36.8 - 41.8%] | 0.102 | 0.12 | NZ_CP064172.1 |
| *V. crassostreae* LMG 22240<sup>T</sup> | 37.8 | [35.4 - 40.3%] | 0.1071 | 0.44 | NZ_AP025476.1 |
| *V. gigantis* ACE001 | 35.8 | [33.4 - 38.3%] | 0.1149 | 0.23 | NZ_CP092384.1 |
| *V. crassostreae* ED395 | 39.4 | [36.9 - 41.9%] | 0.1015 | 0.31 | NZ_CP064170.1 |
| *V. cyclitrophicus* ED287 | 39.3 | [36.8 - 41.8%] | 0.1018 | 0.01 | NZ_CP065366.1 |
| *V. gigantis* LMG 22741<sup>T</sup> | 35.7 | [33.3 - 38.3%] | 0.1151 | 0.25 | NZ_AP025492.1 |
| *V. splendidus* LMG 19031<sup>T</sup> | 43.6 | [41.1 - 46.2%] | 0.0881 | 0.15 | NZ_AP025508.1 |
| *V. pomeroyi* LMG 20537<sup>T</sup> | 34.9 | [32.5 - 37.4%] | 0.1185 | 0.46 | NZ_AP025506.1 |
| *V. lentus* LMG 21034<sup>T</sup> | 38.6 | [36.1 - 41.1%] | 0.1043 | 0.09 | NZ_AP025499.1 |
| *V. chagasii* LMG 21353<sup>T</sup> | 30.4 | [28 - 32.9%] | 0.1402 | 0.41 | NZ_AP025465.1 |
| *V. artabrorum* CECT 7226<sup>T</sup> | 31.8 | [29.4 - 34.3%] | 0.1327 | 0.16 | NZ_AP025458.1 |
| *V. gallaecicus* CECT 7244<sup>T</sup> | 24.4 | [22.1 - 26.8%] | 0.179 | 2.44 | NZ_AP025490.1 |
| *V. cholerae* El Tor N16961 | 20.9 | [18.7 - 23.3%] | 0.2103 | 3.5 | NC_002505 |
| *V. celticus* 7224<sup>T</sup> | 38.5 | [36 - 41%] | 0.1047 | 0.48 | NZ_MVJF01000005.1 |

**Figure 2.**
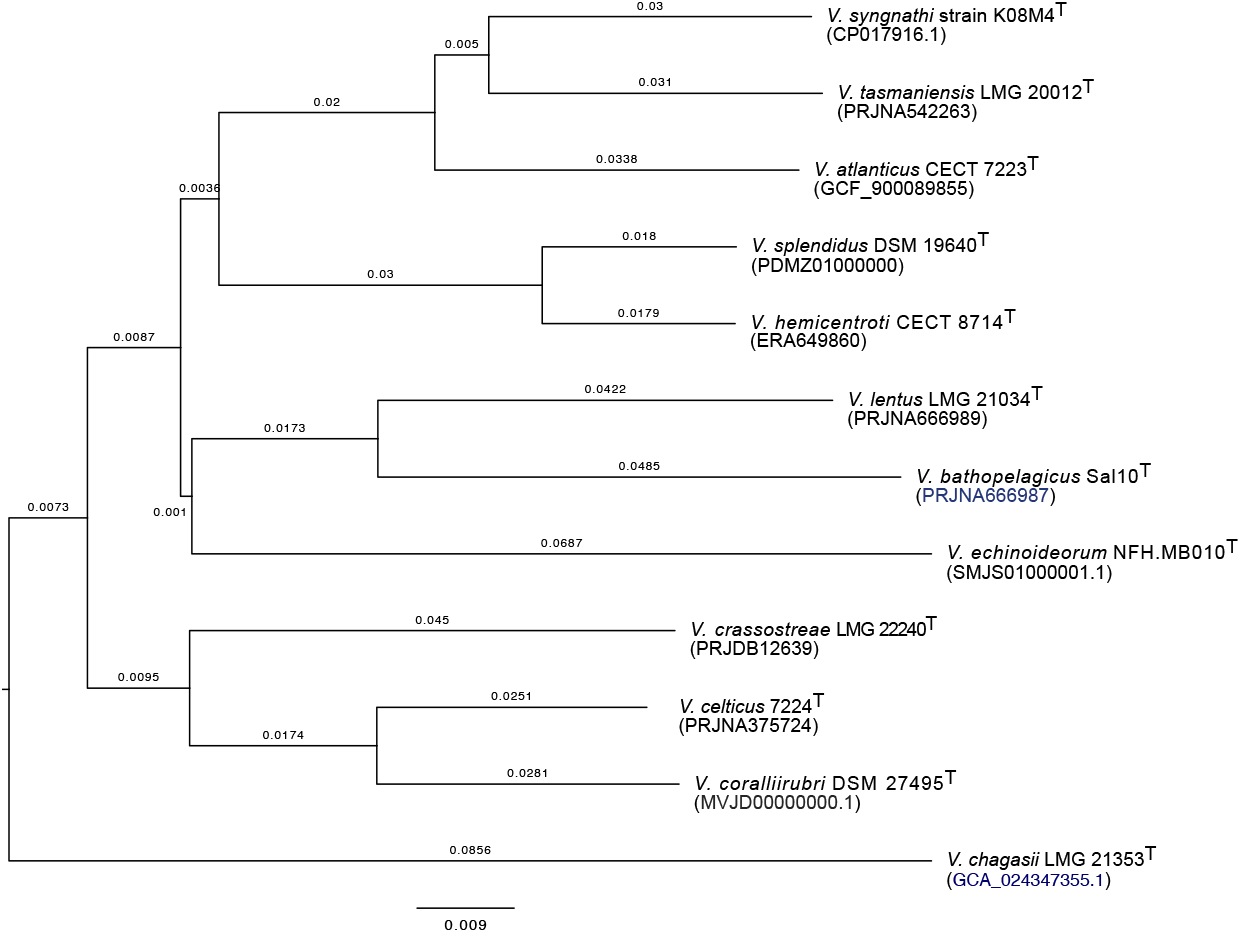
Phylogenetic tree inferred with FastME 2.1.6.1 [34] from GBDP distances calculated from genome sequences. The branch lengths are scaled in terms of GBDP distance formula *d5*. The numbers above branches are GBDP pseudo-bootstrap support values > 60 % from 100 replications, with an average branch support of 97.7 %. The tree was rooted at the midpoint [35].

In addition, phylogenetic analysis based on an MLSA tree using eight housekeeping genes (*ftsZ, gapA, gyrB, mreB, pyrH, recA, rpoA*, and *tpoA*) confirmed that strain K08M4^T^ forms a monophyletic group within the *V. Splendidus* clade (Figure S1).

Furthermore, we compared the genomic relatedness based on the average nucleotide identity (ANI) between strain K08M4^T^ and all closed *Vibrio* sequences as reference available on NCBI (date of access 17.11.2022) using FastANI (https://github.com/ParBLiSS/FastANI; [37]). All closed *Vibrio* genomes were downloaded using the NCBI Datasets command line tools and the following command line “datasets download genome taxon 662 --complete”. Firstly, the ANI values between the complete genome sequence of strain K08M4^T^, were computed separately for Chromosomes1 and Chromosome 2, as well as for the concatenated sequences of Chromosome 1 and Chromosome 2 of all deduplicated closed *Vibrio* genomes (n=532). ANI between the K08M4^T^ comparisons and reference strains that were above 90% from the concatenated sequences were selected for the purpose of visualization (Figure 3) using PyANI [38] and the ANIb option (https://github.com/widdowquinn/pyani). Most ANI values in any combination were below the threshold of 95%, the repeatedly established species demarcation cut-off value [37]. One exception is nucleotide identity between K08M4^T^ strain and *V. tasmaniensis* LMG 20012 (ANI chromosome 1 = 95.5%, chromosome 2 = 93.96%, both chromosomes together = 95.03%).

**Figure 3:**
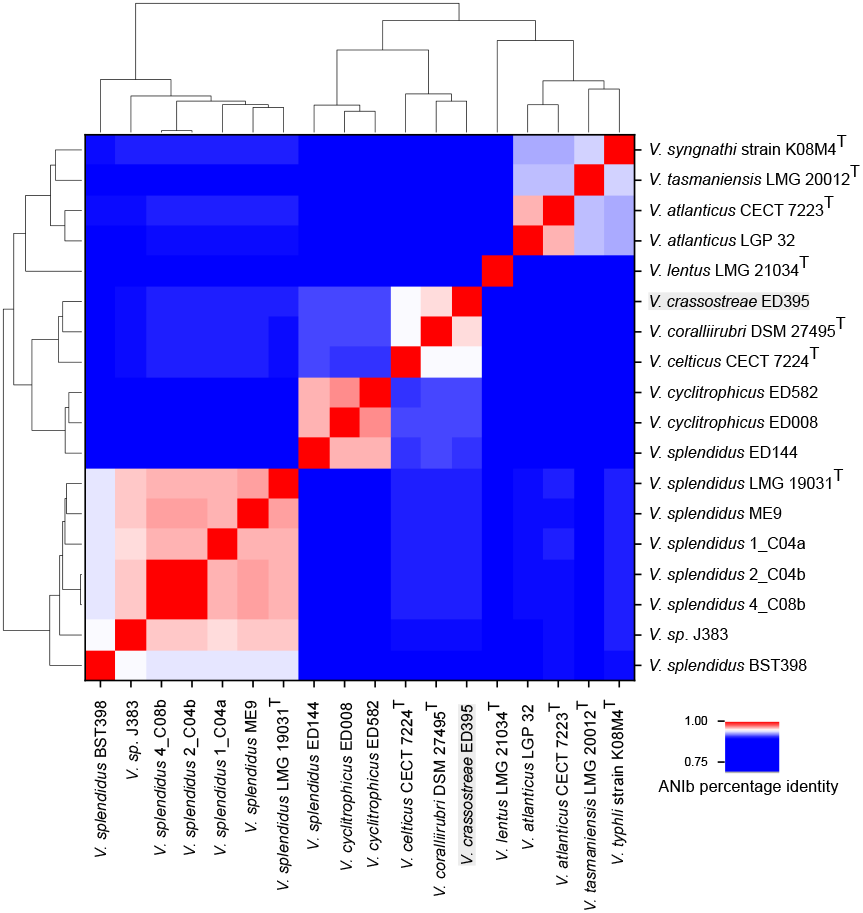
Average nucleotide identity (ANI) analysis of strain K08M4^T^ sequenced strain with available closed *Vibrio* genomes (ANI > 90%) as references. ANI analysis based on BLAST alignment of the genome sequences was performed and visualized using PyANI.

Phylogenomic treeing and taxonomy assignment of K08M4^T^ was additionally performed based on 120 bacterial marker genes with GTDB-Tk v2.1.0 [39, 40] (Database release R207) using the ‘classify_wf’ function and default parameters. Genomes are assigned at the species level if the ANI to the closest GTDB species representative genome was ≥95% and the AF was ≥30%. Accordingly, K08M4^T^ was characterized as *V. tasmaniensis* based on GTDB taxonomy. However, taking all sequence-based results (ANI of K08M4^T^ and *V. tasmaniensis* LMG 20012 is 95.03%, directly at the boundary of the species demarcation cut-off) together with experimental confirmation, we can confidently confirm that K08M4^T^ represent a novel *Vibrio* species.

### Pathogenicity

We performed a controlled infection experiment to test the virulence of strain K08M4^T^ on pipefish larvae (a detailed description of the methods and the statistical analysis can be found in [19]). Briefly, we exposed 10-11 juvenile pipefish in triplicates to either 10^9^ CFU/ml of strain K08M4^T^, 10^9^ CFU/ml of a non-virulent *V. alginolyticus* K10K4, or seawater as control. Fish mortality was recorded daily. On day three, we sampled one fish per experimental unit and analyzed the expression of 32 immune genes relative to two housekeeping genes using a Fluidigm BioMark™ as described in [41]. This included 17 target genes assigned to the innate immune system, two target genes assigned to the complement component system, and 13 target genes assigned to the adaptive immune system (Table S5). Details about function of genes, sequences and primer design can be found in [41].

Strain K08M4^T^ causes high rates of mortality (Figure 4a) in pipefish larvae and elicits a different expression profile of selected immune genes compared to the non-virulent *V. alginolyticus* strain K10K4 or pipefish exposed to seawater (Figure 4b).

**Figure 4.**
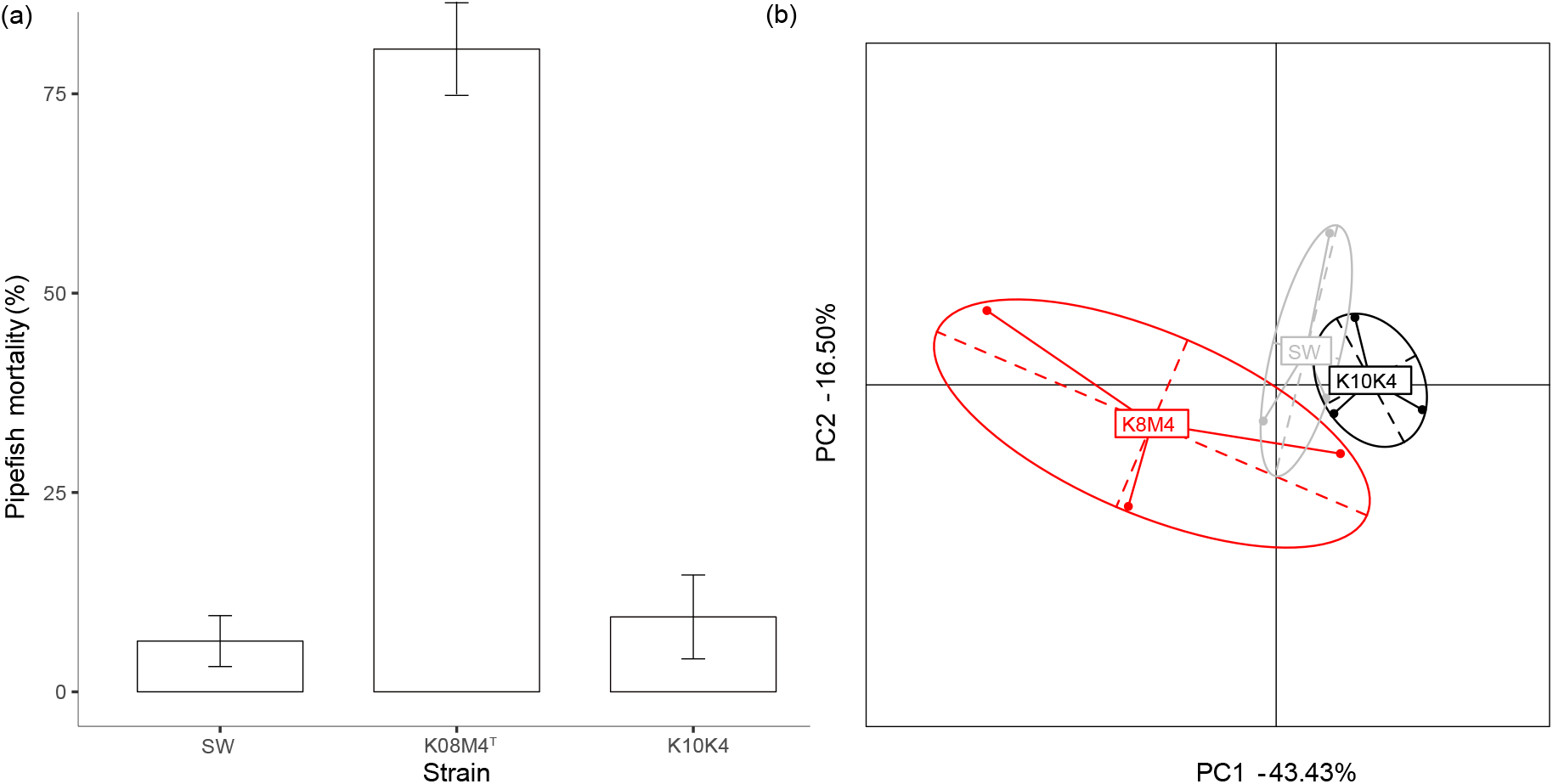
Pipefish mortality in per cent (a) and ordination of differentially expressed immune genes between K08M4, K10K4 or seawater-injected pipefish (SW).

Several genetically encoded virulence factors such as iron transport systems, flagellum/ motility, hemolysins, proteases, lipopolysaccharides, exopolysaccharides, repeats in toxins (RTX), outer membrane proteins, and a type IV pilus could be identified (Table S6). Compared to other *Vibrio* species isolated from the Kiel Fjord, K08M4^T^ has a unique virulence profile [42].

Based on the comparative genomic analysis and phenotypic description, strain K08M4^T^ represents a novel species of the genus *Vibrio*, with an increased virulence profile for juvenile pipefish, for which we propose the name *Vibrio syngnathi* sp. nov.

### Description of *Vibrio syngnathi* sp. nov

*Vibrio syngnathi* (syn.gna’thi. N.L. gen. n. syngnathi, pertaining to the broad-nosed pipefish *Syngnathus typhle* from which the strain was originally isolated)

Cells are slightly curved rods, Gram-stain-negative, curved rod-shaped and motile by means of a single polar flagellum. Colonies are circular, yellow, and 2-4 mm in size after 24h growth on TCBS agar at 25° C. Cells can grow from 9-40° C and pH 4 - 10.5. Optimal growth is observed at 25 - 35° C. Growth occurs in the presence of 0.4 - 12 % (w/v) NaCl. Biochemical characteristics of K08M4^T^ include positive for oxidase, catalase, chitinase, indole production and O1/129 sensitivity. Acid is produced from glycerol, ribose, glucose, fructose, mannose, N-acetylglucosamine, esculin, salicin, cellobiose, maltose, sucrose, trehalose, starch, glycogen, and gluconate. K08M4^T^ is negative for β-galactosidase, arginine dihydrolase, lysing decarboxylase, ornithine decarboxylase, citrate utilization, H2S production, urease, tryptophan deaminase, Voges-Proskauer test, gelatin hydrolysis. The type-strain can grow on glycerol, starch, and glycogen. The major fatty acid (> 10%) are C_16:1_ *ω*7*c* and C_16:0_. K08M4^T^ causes mortality in juvenile pipefish, *S. typhli*.

The type strain, K08M4^T^ (= DSM 109818^T^ = CECT 30086^T^), was isolated from the whole intestines of an adult broad-nosed pipefish *Syngnathus typhle*, caught in the Kiel Fjord, Germany. The DNA G+C content of the type-strain is 44.06%. The genome is deposited at ENA (PRJNA345286) and NCBI (CP017916 and CP017917). The 16S rRNA sequence can be found at NCBI GenBank under the accession number OP359305.

## Abbreviations

TCBS: thiosulfate-citrate-bile-sucrose;
TYGS: Type Strain Genome Server,
dDDH: digital DNA-DNA hybridization

## Funding

This study was supported by the Deutsche Forschungsgemeinschaft (DFG) SPP 1819 on rapid adaptation (WE5822/1-1, WE5822/1-2, and OR4628/4-2 given to CCW and OR) and a Postdoctoral fellowship from the Cluster of Excellence ‘The Future Ocean’ given to CCW.

## Ethical approval

Approval for using pipefish during infection experiments was given by the Ministerium für Landwirtschaft, Umwelt und ländliche Räume des Landes Schleswig-Holstein V312-7224.121-19 (65-5/13).

## Conflict of interests

We declare no conflict of interests.

## Authors’ contribution

OR isolated the strain. CCW, OR and HL designed the study. CCW, RH, HL, BB, CS and JO were involved in the preparation of bacterial DNA and genome sequencing. CCW performed infection experiments. HG, KSW MM, and MNS performed microbiological lab work. CC, HL and BB analysed the bacteria genomes. MH performed electron microscopy. CCW coordinated the project. CCW and CC wrote the first draft of the manuscript. All authors approved the final version of the manuscript.

## Acknowledgements

We are grateful to Anja Frühling, Gesa Martens, Anika Wasner, Bettina Ehlert, Simone Schrader and Nicole Heyer for excellent technical assistance.

## Notes

### Competing Interest Statement

The authors have declared no competing interest.

