## supplementary material for "COMPLETE GENOME SEQUENCE OF *VIBRIO SYNGNATHI SP. NOV*., A FISH PATHOGEN, ISOLATED FROM THE KIEL FJORD"

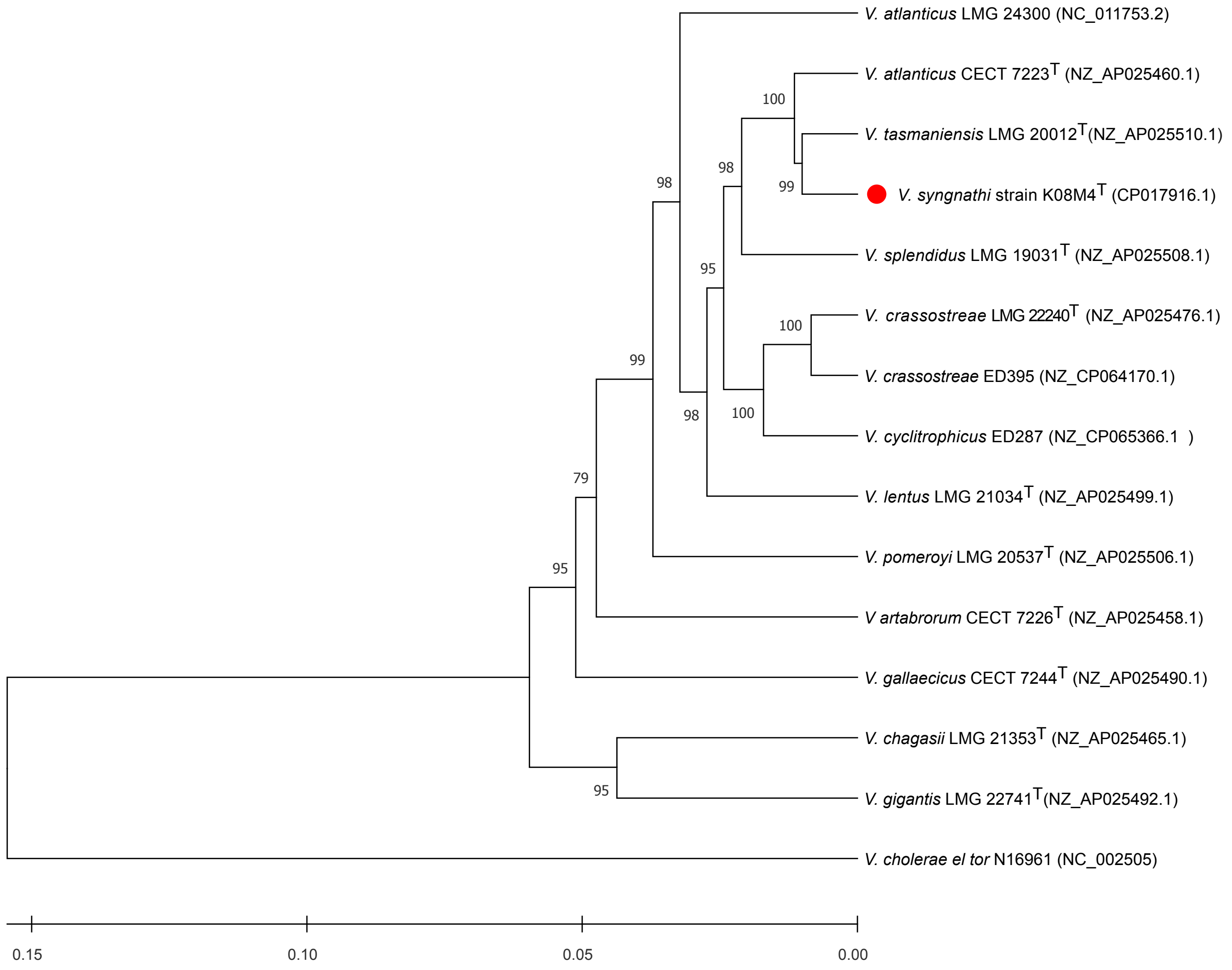


**Figure S1 Phylogenetic tree based on MLSA using eight housekeeping genes** (ftsZ, gapA, gyrB, mreB, pyrH, recA, rpoA, and tpoA**.** The evolutionary history was inferred using the UPGMA method [1]. The optimal tree is shown. The percentage of replicate trees in which the associated taxa clustered together in the bootstrap test (500 replicates) are shown next to the branches [2]. The tree is drawn to scale, with branch lengths in the same units as those of the evolutionary distances used to infer the phylogenetic tree. The evolutionary distances were computed using the Maximum Composite Likelihood method [3] and are in the units of the number of base substitutions per site. This analysis involved 15 nucleotide sequences. There were a total of 11681 positions in the final dataset. Evolutionary analyses were conducted in MEGA11 [4].

**Table S1**: Results from the API ZYM
Strains: 1. DSM 109818$^T$ (this study), 2. *V. tasmaniensis* DSM 17182$^T$ (this study), 3. *V. splendidus* DSM 19640$^T$ (this study), 4 *V. splendidus* DSM 26178$^T$ (this study).

| | 1 | 2 | 3 | 4 |
|---|---|---|---|---|
| Alcaline Phosphatase | 5 | 5 | 5 | 5 |
| Esterase | 3 | 3 | 3 | 2 |
| Esterase Lipase | 3 | 3 | 2 | 1 |
| Lipase | 0 | 0 | 0 | 0 |
| Leucin-Arylamidase | 3 | 3 | 4 | 3 |
| Valin-Arylamidase | 0.5 | 0.5 | 1 | 0.5 |
| Cystin-Arylamidase | 0 | 0 | 0 | 0 |
| Trypsin | 1 | 1 | 2 | 4 |
| Chymotrypsin | 0 | 0 | 0 | 0 |
| Acid Phosphatase | 4 | 5 | 5 | 5 |
| Naphtol-AS-BI-Phosphohydrolase | 2 | 3 | 3 | 3 |
| α-Galactosidase | 0 | 0 | 0 | 0 |
| β-Galactosidase | 0 | 0 | 0 | 0 |
| β-Glucuronidase | 0 | 0 | 0 | 0 |
| α-Glucosidase | 0 | 0 | 0 | 0 |
| β-Glucosidase | 0 | 0 | 0 | 0 |
| N-Acetyl-β-Glucosaminidase | 0 | 0 | 0 | 0 |
| α-Mannosidase | 0 | 0 | 0 | 0 |
| α-Fucosidase | 0 | 0 | 0 | 0 |

**Table S2**: Results from API 20E
Strains: 1. DSM 109818$^T$ (this study), 2. *V. tasmaniensis* DSM 17182$^T$ (this study), 3. *V. splendidus* DSM 19640$^T$ (this study), 4 *V. splendidus* DSM 26178$^T$ (this study).

| | 1 | 2 | 3 | 4 |
|---|---|---|---|---|
| β-Galactosidase (ONPG) | - | - | + | + |
| Arginine dihydrolase (ADH) | - | + | + | + |
| Lysine decarboxylase (LDC) | - | - | - | - |
| Ornithine decarboxylase (ODC) | - | - | - | - |
| Citrate utilization | - | - | - | - |
| $H_2S$ production | - | - | - | - |
| Urease | - | - | - | - |
| Tryptophan deaminase | - | - | - | - |
| Indole | + | + | + | + |
| Voges-Proskauer (VP) | - | - | - | - |
| Gelatine hydrolysis | - | - | + | + |
| Acid from | | | | |
| Glucose | + | + | + | + |
| Mannose | - | + | + | + |
| Inositol | - | - | - | - |
| Sorbitol | - | - | - | - |
| Rhamnose | - | - | - | - |
| Sucrose | + | - | - | - |
| Melibiose | - | - | + | - |
| Amygdalin | + | + | + | + |
| Arabinose | - | - | - | - |

**Table S3**: Results from API 50 CHE; Acid production. Strains: 1. DSM 109818$^T$ (this study), 2. *V. tasmaniensis* DSM 17182$^T$ (this study), 3. *V. splendidus* DSM 19640$^T$ (this study), 4 *V. splendidus* DSM 26178$^T$ (this study).

| | 1 | 2 | 3 | 4 |
|---|---|---|---|---|
| Glycerol | + | + | + | + |
| Erythritol | - | - | - | - |
| D-Arabinose | - | - | - | - |
| L-Arabinose | - | - | - | - |
| Ribose | + | + | + | + |
| D-Xylose | - | - | - | - |
| L-Xylose | - | - | - | - |
| Adonitol | - | - | - | - |
| βMeDXyloside | - | - | - | - |
| Galactose | - | - | + | + |
| Glucose | + | + | + | + |
| Fructose | + | + | + | + |
| Mannose | + | + | + | + |
| Sorbose | - | - | - | - |
| Rhamnose | - | - | - | - |
| Inositol | - | - | - | - |
| Mannitol | - | + | + | + |
| Sorbitol | - | - | - | - |
| αMeDMannoside | - | - | - | - |
| αMeGlucoside | - | - | - | - |
| N-Acetylglucosamine | + | + | + | + |
| Amygdalin | - | - | + | - |
| Arbutin | - | - | - | - |
| Esculin | + | + | + | + |
| Salicin | + | + | + | - |
| Cellobiose | + | + | + | + |
| Maltose | + | + | + | + |
| Lactose | - | - | - | + |
| Melibiose | - | - | + | - |
| Sucrose | + | - | - | - |
| Trehalose | + | + | + | + |
| Inulin | - | - | - | + |
| Melizitose | - | - | - | - |
| Raffinose | - | - | - | - |
| Starch | + | + | + | + |
| Glycogen | + | + | + | + |
| Xylitol | - | - | - | - |
| Gentibiose | - | - | + | - |
| D-Turanose | - | - | - | - |
| D-Lyxose | - | - | - | - |
| D-Tagatose | - | - | - | - |
| D-Fucose | - | - | - | - |
| L-Fucose | - | - | - | - |
| D-Arabitol | - | - | - | - |
| L-Arabitol | - | - | - | - |
| Gluconate | + | + | + | + |
| 2Ketogluconate | - | - | + | - |
| 5Ketogluconate | - | ? | - | - |

**Table S4:** Fatty acids for strain K08M4$^{T}$

| MIDI peak naming/TSBA 6 | GC/MS | % |
|---|---|---:|
| 10:00 | 10:00 | <0.1 |
| 10:0 3OH | 10:0 3OH | <0.1 |
| 12:0 iso | 12:0 iso | 0.1 |
| unknown 11.799 | Identified as 12:1 w7c | 0.2 |
| 12:0 | 12:0 | 3.9 |
| 13:0 | 13:0 | <0.1 |
| 12:0 iso 3OH | 12:0 iso 3OH | 0.2 |
| 12:0 3OH | 12:0 3OH | 2.7 |
| 14:0 iso | 14:0 iso | 0.8 |
| -- | Identified as 14:1 w7c | 0.6 |
| 14:1 w5c | 14:1 w5c | <0.1 |
| 14:0 | 14:0 | 3.8 |
| Summed Feature 1 (13:0 3OH/15:1 i H) | Identified as 13:0 3OH | 0.1 |
| 15:1 w8c | 15:1 w8c | 0.2 |
| 15:0 | 15:0 | 0.9 |
| 14:0 iso 3OH | 14:0 iso 3OH | 0.7 |
| Summed Feature 2 (14:0 3OH/16:1 iso I) | Identified as 14:0 3OH | 1.8 |
| 16:0 iso | 16:0 iso | 7.6 |
| Summed Feature 3 (16:1 w7c/16:1 w6c) | Identified as 16:1 w7c | 39.9 |
| Summed Feature 3 (16:1 w6c/16:1 w7c) | Identified as 16:1 w7t | 3.3 |
| 16:1 w5c | 16:1 w5c | 0.2 |
| 16:0 | 16:0 | 19.2 |
| 17:1 w8c | 17:1 w8c | 0.5 |
| 17:1 w6c | 17:1 w6c | 0.2 |
| 17:0 | 17:0 | 0.4 |
| 16:0 iso 3OH | 16:0 iso 3OH | <0.1 |
| 16:0 3OH | 16:0 3OH | 0.2 |
| 18:0 iso | 18:0 iso | 0.9 |
| Summed Feature 8 (18:1 w7c/18:1 w6c) | Identified as 18:1 w7c | 8.7 |
| 18:1 w5c | 18:1 w5c | 0.2 |
| 18:0 | 18:0 | 1.0 |
| 18:1 w7c 11-methyl | 18:1 w7c 11-methyl | 0.2 |
| Summed Feature 7 (un 18.846/19:1 w6c) | Identified as 10,13-epoxy-11-methyl- octadecadienoate | 0.7 |
| 20:0 iso | 20:0 iso | <0.1 |
| 20:1 w7c | 20:1 w7c | 0.5 |

**Table S5**: Genes analysed from the infection experiment and their classification as either: adaptive immune gene, complement system, or innate immune system.

| Gene | Type |
|---|---|
| bcell_rap31 | adaptive |
| CD45 | adaptive |
| CD74 | adaptive |
| Fut9 | adaptive |
| H2K1 | adaptive |
| HIVEP2 | adaptive |
| HIVEP3 | adaptive |
| IgM | adaptive |
| integ | adaptive |
| lymphag75 | adaptive |
| lymphcyt | adaptive |
| MEF2C | adaptive |
| tap | adaptive |
| C3 | complement |
| C9 | complement |
| AIF | innate |
| calrucul | innate |
| cf | innate |
| ck7 | innate |
| hsp60 | innate |
| il10 | innate |
| il8 | innate |
| intf | innate |
| kin | innate |
| lectpt1 | innate |
| lectpt2 | innate |
| lps_tnf | innate |
| nramp | innate |
| tnf | innate |
| tranfe | innate |
| tspo | innate |
| tyroprot | innate |

**Table S6** Genetically encoded virulence factors of *Vibrio syngnathi* K08M4$^T$

| Function | Gene id | Annotation |
|---|---|---|
| Iron transport systems | K08M4_02090 | fief, Ferrous-iron efflux pump |
| | K08M4_04510 | fhuC_1, Iron(3+)-hydroxamate import TP-binding protein |
| | K08M4_05830 | futA1, Iron uptake protein A1 precursor |
| | K08M4_20140 | fhuC_2, Iron(3+)-hydroxamate import ATP-binding protein FhuC |
| | K08M4_20150 | fhuD_1, Iron(3+)-hydroxamate-binding protein FhuD |
| | K08M4_20160 | fhuB_1, Iron(3+)-hydroxamate import system permease protein FhuB |
| | K08M4_21150 | feoB, Ferrous iron transport protein B |
| | K08M4_21160 | feoA, Ferrous iron transport protein A |
| | K08M4_34650 | fhuC_3, Iron(3+)-hydroxamate import ATP-binding protein FhuC |
| | K08M4_34680 | fhuA_1, Ferrichrome-iron receptor precursor |
| | K08M4_34690 | fhuD_2, Iron(3+)-hydroxamate-binding protein FhuD |
| | K08M4_34700 | fhuB_2, Iron(3+)-hydroxamate import system permease protein FhuB |
| | K08M4_39310 | fhuC_4, Iron(3+)-hydroxamate import ATP-binding protein FhuC |
| | K08M4_39330 | feuB, Iron-uptake system permease protein FeuB |
| Motility | K08M4_01150 | flagellar basal body-associated protein FliL-like protein |
| | K08M4_08020 | flagellar basal body P-ring biosynthesis protein FlgA |
| | K08M4_08050 | flgB, Flagellar basal body rod protein FlgB |
| | K08M4_08060 | flgC, Flagellar basal-body rod protein FlgC |
| | K08M4_08080 | flgE, Flagellar hook protein FlgE |
| | K08M4_08090 | flgF, Flagellar basal-body rod protein FlgF |
| | K08M4_08100 | flgG, Flagellar basal-body rod protein FlgG |
| | K08M4_08110 | flgH, Flagellar L-ring protein precursor |
| | K08M4_08120 | flgI, Flagellar P-ring protein precursor |
| | K08M4_08140 | flgK, Flagellar hook-associated protein 1 |
| | K08M4_08150 | flgL, Flagellar hook-associated protein 3 |
| | K08M4_08160 | flaB_1, Flagellin B |
| | K08M4_08180 | flaD_1, Flagellin D |
| | K08M4_08190 | flaB_2, Flagellin B |
| | K08M4_08200 | flagellar protein FlaG |
| | K08M4_08210 | fliD, Flagellar hook-associated protein 2 |
| | K08M4_08220 | Flagellar protein FliT |
| | K08M4_08230 | fliS, Flagellar protein FliS |
| | K08M4_08270 | fliE, Flagellar hook-basal body complex protein FliE |
| | K08M4_08280 | fliF, Flagellar M-ring protein |
| | K08M4_08290 | fliG, Flagellar motor switch protein FliG |
| | K08M4_08300 | fliH, Flagellar assembly protein FliH |
| | K08M4_08310 | fliI, Flagellum-specific ATP synthase |
| | K08M4_08320 | flagellar biosynthesis chaperone |
| | K08M4_08330 | Flagellar hook-length control protein FliK |
| | K08M4_08340 | flagellar basal body-associated protein FliL |
| | K08M4_08350 | fliM, Flagellar motor switch protein FliM |
| | K08M4_08360 | fliN, Flagellar motor switch protein FliN |
| | K08M4_08370 | fliO, Flagellar protein FliO |
| | K08M4_08380 | fliP, Flagellar biosynthetic protein FliP precursor |

| | | |
|---|---|---|
| | K08M4_08390 | fliQ, Flagellar biosynthetic protein FliQ |
| | K08M4_08400 | flagellar biosynthesis protein FliR |
| | K08M4_08410 | flhB, Flagellar biosynthetic protein FlhB |
| | K08M4_08420 | flhA, Flagellar biosynthesis protein FlhA |
| | K08M4_08430 | flhF, Flagellar biosynthesis protein FlhF |
| | K08M4_08440 | ylxH, Flagellum site-determining protein YlxH |
| | K08M4_09230 | Flagellin N-methylase |
| | K08M4_14080 | Flagellin N-methylase |
| | K08M4_21920 | flaD_2, Flagellin D |
| | K08M4_21930 | flaD_3, Flagellin D |
| | K08M4_33290 | Flagellar protein YcgR |
| | K08M4_35140 | Flagellin N-methylase |
| | K08M4_36060 | flagellar basal body rod modification protein |
| | K08M4_36470 | flagellar basal body rod modification protein |
| **Hemolysins** | K08M4_01690 | tlyC, Hemolysin C |
| | K08M4_12340 | hlyA, Hemolysin, chromosomal |
| | K08M4_29570 | hemolysin-III related |
| | K08M4_31020 | hlyD_1, Hemolysin secretion protein D, chromosomal |
| | K08M4_31030 | hlyB_1, Alpha-hemolysin translocation ATP-binding protein HlyB |
| | K08M4_31050 | hlyC, Hemolysin-activating lysine-acyltransferase HlyC |
| | K08M4_31830 | Thermolabile hemolysin precursor |
| | K08M4_34780 | hlyD_2, Hemolysin secretion protein D, chromosomal |
| | K08M4_38260 | hlyD_3, Hemolysin secretion protein D, chromosomal |
| | K08M4_38290 | hlyB_2, Alpha-hemolysin translocation ATP-binding protein HlyB |
| **Proteases** | K08M4_01140 | glpG, Rhomboid protease GlpG |
| | K08M4_02200 | ATP-dependent Clp protease ATP-binding subunit ClpX |
| | K08M4_02210 | hslV, ATP-dependent protease subunit HslV |
| | K08M4_06120 | ftsH, ATP-dependent zinc metalloprotease FtsH |
| | K08M4_06290 | yhbU_1, putative protease YhbU precursor |
| | K08M4_08700 | ptrA_1, Protease 3 precursor |
| | K08M4_10590 | ptrB, Protease 2 |
| | K08M4_10730 | clpS, ATP-dependent Clp protease adapter protein ClpS |
| | K08M4_10740 | clpA, ATP-dependent Clp protease ATP-binding subunit ClpA |
| | K08M4_12710 | ypwA, Putative metalloprotease YpwA |
| | K08M4_14380 | yhbU_2, putative protease YhbU precursor |
| | K08M4_15570 | lon_1, Lon protease |
| | K08M4_15700 | prc, Tail-specific protease precursor |
| | K08M4_19120 | htpX_1, Protease HtpX |
| | K08M4_19440 | htpX_2, Protease HtpX |
| | K08M4_20820 | intramembrane serine protease GlpG |
| | K08M4_20860 | lon_2, Lon protease |
| | K08M4_20870 | ATP-dependent Clp protease ATP-binding subunit ClpX |
| | K08M4_20880 | ATP-dependent Clp protease proteolytic subunit |
| | K08M4_21610 | hflK_1, Modulator of FtsH protease HflK |
| | K08M4_22460 | RIP metalloprotease RseP |

| | | |
|---|---|---|
| | K08M4_24160 | yhbU_3, putative protease YhbU precursor |
| | K08M4_25410 | RIP metalloprotease RseP |
| | K08M4_25420 | degS, Serine endoprotease DegS |
| | K08M4_26170 | protease TldD |
| | K08M4_27000 | hflC_1, Modulator of FtsH protease HflC |
| | K08M4_27010 | hflK_2, Modulator of FtsH protease HflK |
| | K08M4_38910 | hflC_2, Modulator of FtsH protease HflC |
| | K08M4_42320 | DNA-binding ATP-dependent protease La |
| | K08M4_43000 | ATP-dependent protease La (LON) domain protein |
| | K08M4_43310 | ptrA_2, Protease 3 precursor |
| | K08M4_43320 | ptrA_3, Protease 3 precursor |
| **Lipopolysaccharide** | K08M4_03560 | lptB, Lipopolysaccharide export system ATP-binding protein LptB |
| | K08M4_03570 | lptA, Lipopolysaccharide export system protein LptA precursor |
| | K08M4_03580 | lptC, Lipopolysaccharide export system protein LptC |
| | K08M4_03820 | lptF, Lipopolysaccharide export system permease protein LptF |
| | K08M4_03830 | lptG, Lipopolysaccharide export system permease protein LptG |
| | K08M4_27870 | rfaQ_1, Lipopolysaccharide core heptosyltransferase RfaQ |
| | K08M4_27880 | rfaQ_2, Lipopolysaccharide core heptosyltransferase RfaQ |
| **repeats in toxins (RTX)** | K08M4_30240 | prsE, Type I secretion system membrane fusion protein PrsE |
| | K08M4_30250 | hypothetical protein |
| | K08M4_30260 | yegE, putative diguanylate cyclase YegE |
| | K08M4_30270 | apxIB_1, Toxin RTX-I translocation ATP-binding protein |
| | K08M4_31010 | apxIB_2, Toxin RTX-I translocation ATP-binding protein |
| | K08M4_31020 | hlyD_1, Hemolysin secretion protein D, chromosomal |
| | K08M4_31030 | hlyB_1, Alpha-hemolysin translocation ATP-binding protein HlyB |
| | K08M4_31040 | hypothetical protein |
| | K08M4_31050 | hlyC, Hemolysin-activating lysine-acyltransferase HlyC |
| | K08M4_31060 | toxA, Dermonecrotic toxin |
| | K08M4_34760 | Toxin RTX-I translocation ATP-binding protein |
| | K08M4_34770 | Toxin RTX-I translocation ATP-binding protein |
| | K08M4_34780 | Hemolysin secretion protein D, chromosomal |
| | K08M4_38260 | Hemolysin secretion protein D, chromosomal |
| | K08M4_38270 | Chemotaxis protein CheY |
| | K08M4_38280 | Outer membrane protein TolC precursor |
| | K08M4_38290 | Alpha-hemolysin translocation ATP-binding protein HlyB |
| | K08M4_38300 | Toxin RTX-I translocation ATP-binding protein |
| **outer membrane proteins** | K08M4_07570 | lolB, Outer-membrane lipoprotein LolB precursor |
| | K08M4_08640 | 47 kDa outer membrane protein precursor |
| | K08M4_13870 | outer membrane protein A |
| | K08M4_14610 | ttgI, Toluene efflux pump outer membrane protein TtgI precursor |
| | K08M4_14830 | oprF_1, Outer membrane porin F precursor |
| | K08M4_15940 | bepC_1, Outer membrane efflux protein BepC precursor |
| | K08M4_15950 | oprF_2, Outer membrane porin F precursor |
| | K08M4_16240 | Outer membrane efflux protein |
| | K08M4_16970 | outer membrane protein A |

| | | |
|---|---|---|
| | K08M4_17430 | bepC_2, Outer membrane efflux protein BepC precursor |
| | K08M4_17440 | oprF_3, Outer membrane porin F precursor |
| | K08M4_19540 | lolA, Outer-membrane lipoprotein carrier protein precursor |
| | K08M4_20110 | blc, Outer membrane lipoprotein Blc precursor |
| | K08M4_21050 | slp, Outer membrane protein slp precursor |
| | K08M4_22080 | bamC, Outer membrane protein assembly factor BamC precursor |
| | K08M4_22450 | bamA, Outer membrane protein assembly factor BamA precursor |
| | K08M4_23500 | bamE, Outer membrane protein assembly factor BamE precursor |
| | K08M4_23720 | bamB, Outer membrane protein assembly factor BamB precursor |
| | K08M4_24300 | bamD, Outer membrane protein assembly factor BamD precursor |
| | K08M4_25310 | outer membrane lipoprotein |
| | K08M4_25490 | tolC_1, Outer membrane protein TolC precursor |
| | K08M4_30000 | ompA_1, Outer membrane protein A precursor |
| | K08M4_31810 | lpp, Major outer membrane lipoprotein Lpp precursor |
| | K08M4_33320 | phoE, Outer membrane pore protein E precursor |
| | K08M4_33640 | ompA_2, Outer membrane protein A precursor |
| | K08M4_34020 | tolC_2, Outer membrane protein TolC precursor |
| | K08M4_34500 | oprF_4, Outer membrane porin F precursor |
| | K08M4_34720 | Putative outer membrane protein precursor |
| | K08M4_34810 | Outer membrane efflux protein |
| | K08M4_37160 | outer membrane-specific lipoprotein transporter subunit LolE |
| | K08M4_37320 | 47 kDa outer membrane protein precursor |
| | K08M4_38280 | tolC_3, Outer membrane protein TolC precursor |
| **Type IV pilus** | K08M4_34370 | Type IV leader peptidase family protein |
| | K08M4_34380 | Flp/Fap pilin component |
| | K08M4_34390 | hypothetical protein |
| | K08M4_34400 | type II secretory pathway protein |
| | K08M4_34410 | hypothetical protein |
| | K08M4_34420 | CobQ/CobB/MinD/ParA nucleotide binding domain protein |
| | K08M4_34430 | P-type DNA transfer ATPase VirB11 |
| | K08M4_34440 | Bacterial type II secretion system protein F domain protein |
| | K08M4_34450 | Bacterial type II secretion system protein F domain protein |
